# Cetacean “gas-bubble thromboembolic polycystic liver disease”: “Budd-Chiari like syndrome” in dolphins?

**DOI:** 10.1101/2022.03.05.483097

**Authors:** Antonio Fernandez, Paul D Jepson, Josue Diaz-Delgado, Yara Bernaldo de Quiros, Eva Sierra, Blanca Mompeo, Ana Isabel Vela, Giovanni Di Guardo, Cristian Suarez-Santana, Antonio Espinosa de los Monteros, Pedro Herraez, Marisa Andrada, Maria Jose Caballero, Miguel Rivero, Francesco Consoli, Ayoze Castro, Oscar Quesada, Manuel Arbelo

## Abstract

Nearly two decades ago, pathologic examination results suggested acoustic factors, such as mid-frequency active naval military sonar (MFAS) could be the cause of acute decompression-like sickness in stranded beaked whales. Acute systemic gas embolism in these beaked whales was published together with enigmatic cystic liver lesions (CLL), characterized by intrahepatic encapsulated gas-filled cysts, tentatively interpreted as “gas-bubble” lesions in various cetacean species. Here we provide a pathologic reinterpretation of CLL in cetaceans. From 1,200 cetaceans necropsied, CLL were only observed in striped dolphins (*Stenella coeruleoalba*), with a low prevalence (2%), and recapitulated pathologic features of Budd-Chiari syndrome in humans. Our results strongly suggest that CLL are the result of the combination of pre-existing or concomitant hepatic vascular disorder (e.g., severe hepatobiliary trematodiasis) superimposed and exacerbated by gas bubbles, and clearly differ from acute systemic gas embolism in stranded beaked whales linked to MFAS.

## Introduction

In 2003, in an article in *Nature*, Jepson *et al*., described systemic gas embolism in stranded beaked whales associated temporally and spatially with naval maneuvers using mid-frequency active sonar (MFAS) and cystic liver lesions (CLL) of unknown etiology in stranded dolphins^1,2^, suggestive of a chronic form of gas-bubble lesion (gas embolism), potentially reminiscent of decompression sickness (DCS; “the bends”) in humans. This account challenged the long-believed immunity of cetaceans to decompression-related side effects through evolution^1^. The main criticism to the initial hypothesis was the lack of comparable CLL in human DCS^3,4^. Furthermore, these CLL clearly differed from acute gas-embolism-associated lesions described in mass-stranded beaked whales in association with MFAS in the Canary Islands, Spain^5^, cited in Jepson *et al*. (2003). Since then, several studies have demonstrated the presence of *in vivo* and *postmortem* intravascular and parenchymal gas bubbles and associated lesions highly consistent with DCS-like in cetaceans^6,7,8,9^.

The etiopathogenetic mechanisms of gas embolism in cetaceans are not fully understood^9^. Specifically, the very rare, chronic CLL observed in few delphinids, namely UK-stranded harbor porpoise (*Phocoena phocoena*), common dolphins (*Delphinus delphis*), Risso’s dolphins (*Grampus griseus*), and Blainville’s beaked whale (*Mesoplodon densirostris*), and a harbor porpoise stranded off Germany^1,2,10^, their potential association with anthropic acoustic sources, as well as their elusive analogy to DCS-like in MFAS-associated beaked whales’ mass strandings remain a subject of controversy and discussion. We hypothesized that those CLL could have a different etiopathogenesis, specifically, one that would involve a pre-existing hepatic vascular disorder e.g., phlebothrombosis and entrapment of decompression-related intravascular gas bubbles, in absence of any demonstrable primary relation with anthropic sonar activities.

## Results

From 1,200 cetaceans and 172 striped dolphins examined, four striped dolphins (cases 1 through 4) exhibited lesions consistent with CLL^1,2^. Age categories of examined animals ranged from subadult (n=1) to adult (n=3). All dolphins were males. Two animals had poor body condition; two animals were emaciated. Three animals were first seen stranded dead; one dolphin (case 4) stranded alive (Table 1). Upon physical examination, case 4 presented extensive rostral maxillary and mandibular fractures with soft tissue loss, had altered buoyancy characterized by lateral deviation and up-side-down position, and kept its eyes closed. Multiple reintroduction attempts were unsuccessful, and the animal died shortly after, within 2.5 hours of initial sighting.

**Table 1.** Biologic and epidemiologic data of striped dolphins included in this study.

| Case | Sex | Length (cm) | Age | Stranding date | Stranding location | Stranding condition | Decomposition code | Body condition |
| --- | --- | --- | --- | --- | --- | --- | --- | --- |
| 1 | Male | 231 | Adult | 9/25/07 | Fuerteventura | Dead | 2 | Emaciated |
| 2 | Male | 206 | Subadult | 5/19/14 | La Graciosa | Dead | 4 | Poor |
| 3 | Male | 215 | Adult | 5/13/16 | Tenerife | Dead | 3 | Emaciated |
| 4 | Male | 234 | Adult | 26/06/17 | Bizkaia | Alive | 1 | Poor |

### Pathologic and immunohistochemical examinations

The macroscopic features of CLL were similar among individuals with variability concerning the extent of the parenchyma involved. Affecting from less than 5% (case 1) up to more than 80% (case 4) of the liver, primarily the right liver lobe, there were numerous, multifocal to coalescing, 2-20 mm in diameter, variably firm, raised nodules. On cut surface, these nodules comprised cavities filled with gas (see gas analyses below) and/or red friable material (fibrin, blood) that expanded the parenchyma (Figure 1). All animals had varying degrees of hepatic sinus and intrahepatic vein obliterative thromboses noticeable as segmental expansions of veins from the hilus and varying depths into the liver parenchyma (Figure 2). These venous thrombi were variably expanded by gas-filled cavities.

**Figure 1.**
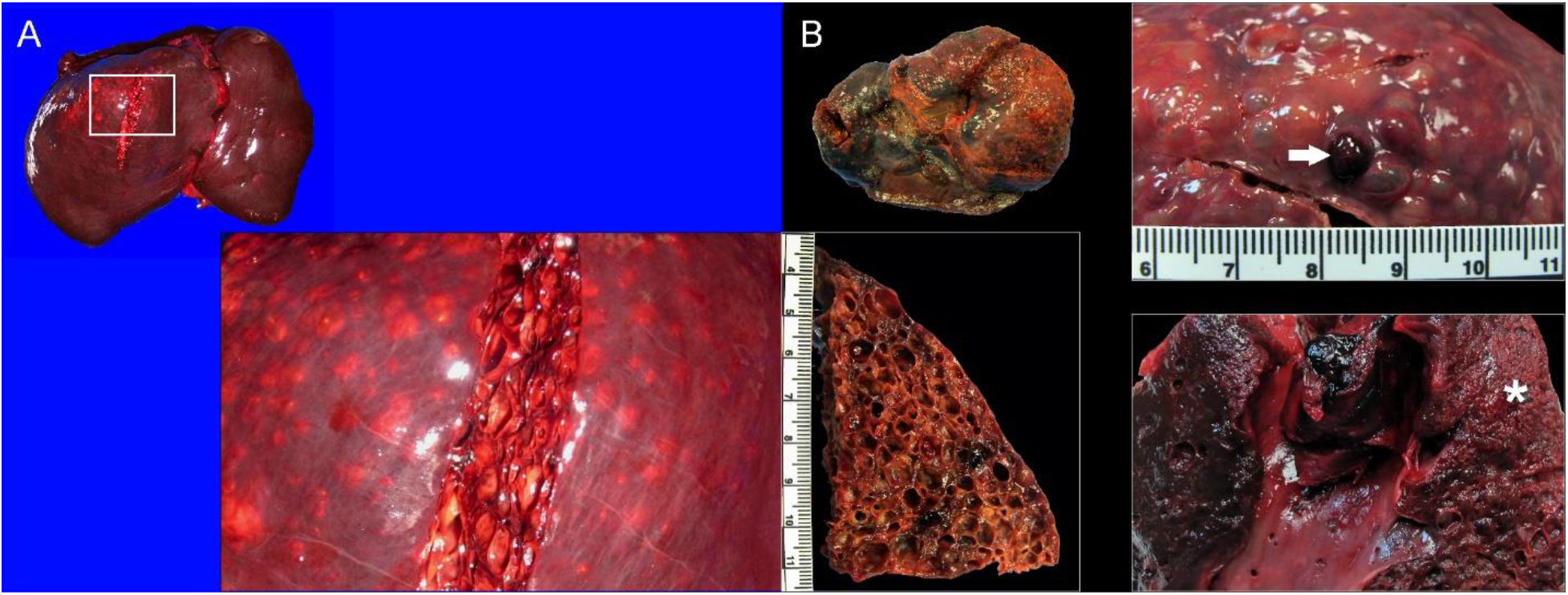
Macroscopic features of cystic liver lesions in dolphins. A) Case 3, liver, cranio-caudal view. The parietal aspect of the right liver lobe is expanded by 0.5 to 2 cm in diameter encapsulated, gas-filled cystic nodules. Inset: detail of cystic nodules on cut surface. B) Case 4, liver, caudo-cranial view (visceral aspect). The entire right liver lobe is expanded by 0.2 to 1 cm in diameter cystic nodules. Right upper inset: detail of surface of cystic nodules, including cavities filled with hemorrhage (arrow). Right lower inset: Obliterative thrombus with the right hepatic venous sinus and major branches. The adjacent liver parenchyma is necrotic (infarct, asterisk). Left lower inset: detail of cystic nodules on cut surface.

**Figure 2.**
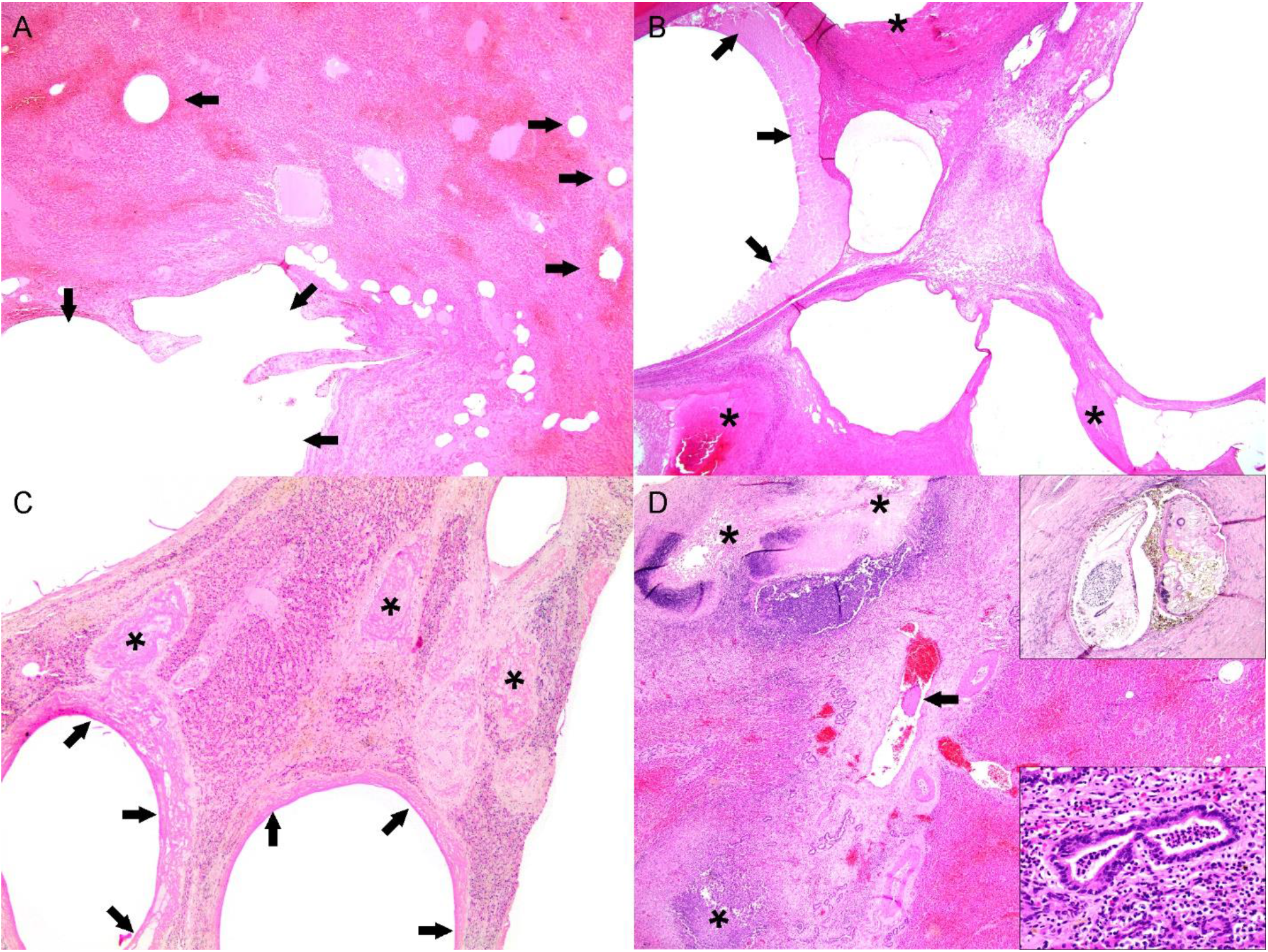
Microscopic features of cystic liver lesions in dolphins. A) Case 4. Acute cystic lesions including cystically dilated veins of varying calibers (arrows) along with hemorrhage and sinusoidal congestion. B) Case 4. Early cystic characterized by negatively-stained (empty, gas-filled) cystic dilatations with serum protein (arrows) and fibrin (asterisks). Remaining hepatic parenchyma exhibits degenerative and atrophic phenomena, as well as hepatocellular loss. C) Case 3. Early to subacute venous cystic dilatations where thick peripheral fibrin (arrows) and thrombi (asterisks) prevail. The adjacent hepatic parenchyma exhibits degenerating and atrophic hepatocytes, fibrin, and mild fibrosis. D) Case 3. Locally extensive pleocellular and fibrosing cholangiohepatitis with venous thrombosis (arrow) and necrosis (asterisk). Right upper inset: intralesional adult trematodes (morphology compatible with *Campula* sp.). Lower inset: suppurative and eosinophilic cholangiohepatitis.

Other relevant gross lesions in these animals were as follows (Supplemental Table 1). Case 1 had severe hepatobiliary infection by *Brachycladiidae* trematodes with multifocal hepatic pyogranulomas, ulcerative trematodal gastritis in the glandular and pyloric gastric compartments, and a large intraperitoneal left peritesticular and epididymal abscess. Case 2 had an 18 cm in diameter, transmural hepato-pyloric abscess that communicated the ulcerated, perforated and fistulized pylorus with the parietal aspect of the right hepatic lobe. On cut surface, there was abundant necrosuppurative exudate with fistulous tracks, numerous adult and larval trematodes compatible with *Campula* sp. and *Pholeter gastrophilus*, varying amounts of ingesta and fish otoliths, and extensive granulation tissue. These lesions resulted in moderate peritoneal effusion and peritonitis.

The histologic features of CLL were similar in all cases with slight variations, primarily relating to their chronicity, and along with their distribution, revealed an ordered progression from hepatic veins into the adjacent hepatic parenchyma. Multifocally, large and medium caliber intrahepatic portal veins had organized obliterative fibrin thrombi with “empty” (gas-filled) cysts (Figure 2). Smaller caliber veins and venules were also consistently affected and had variably cystically distended segments with eosinophilic homogeneous material (serum protein), peripheralized erythrocytes, and fibrin strands. Similar cystic spaces expanded and disrupted the hepatic parenchyma, including some cysts that were lined by few endothelial cells, and dilated sinusoids (peliosis, telangiectasia). Also, there was evidence of hepatocellular pressure and coagulative necrosis, and hepatocellular atrophy. Fibrin was common within the sinusoids and hepatic veins (Figures 2 and 3). There was a clear progression from cystic spaces partially lined or circumferentially lined by fibrin (“acute”) and erythrocytes towards cystic spaces bound by variably dense bands of connective tissue (“chronic”). The latter were mostly composed of collagen, as highlighted by trichrome (Figure 3) and Picrosirius red stains. In chronic lesions, between cysts’ walls, there were collapsed and distorted entrapped portal elements (lymphatics, blood vessels, bile ducts), as well as Kupffer cells laden with hemosiderin and/or hematoidin; the hepatic parenchyma between the cysts was congested and hepatocellular atrophy was consistent. There was no evidence of vascular recanalization in any of the cystic lesions observed yet preserved veins and arteries often exhibited varying degrees of perivascular intimal and mural fibroelastosis, arteriosclerosis and tunica media hypertrophy and hyperplasia. Neovascularization and hepatocellular nodular hyperplasia were not apparent. All these animals had localized to multifocal, moderate to severe trematodal cholangiohepatitis with necrosis (Figure 2D), bile duct hyperplasia, fibrosis, hemorrhage, and hemosiderosis. Cases 1 and 2 had concomitant hepatic intralesional bacteria. Case 3 also had marked cholestasis.

**Figure 3.**
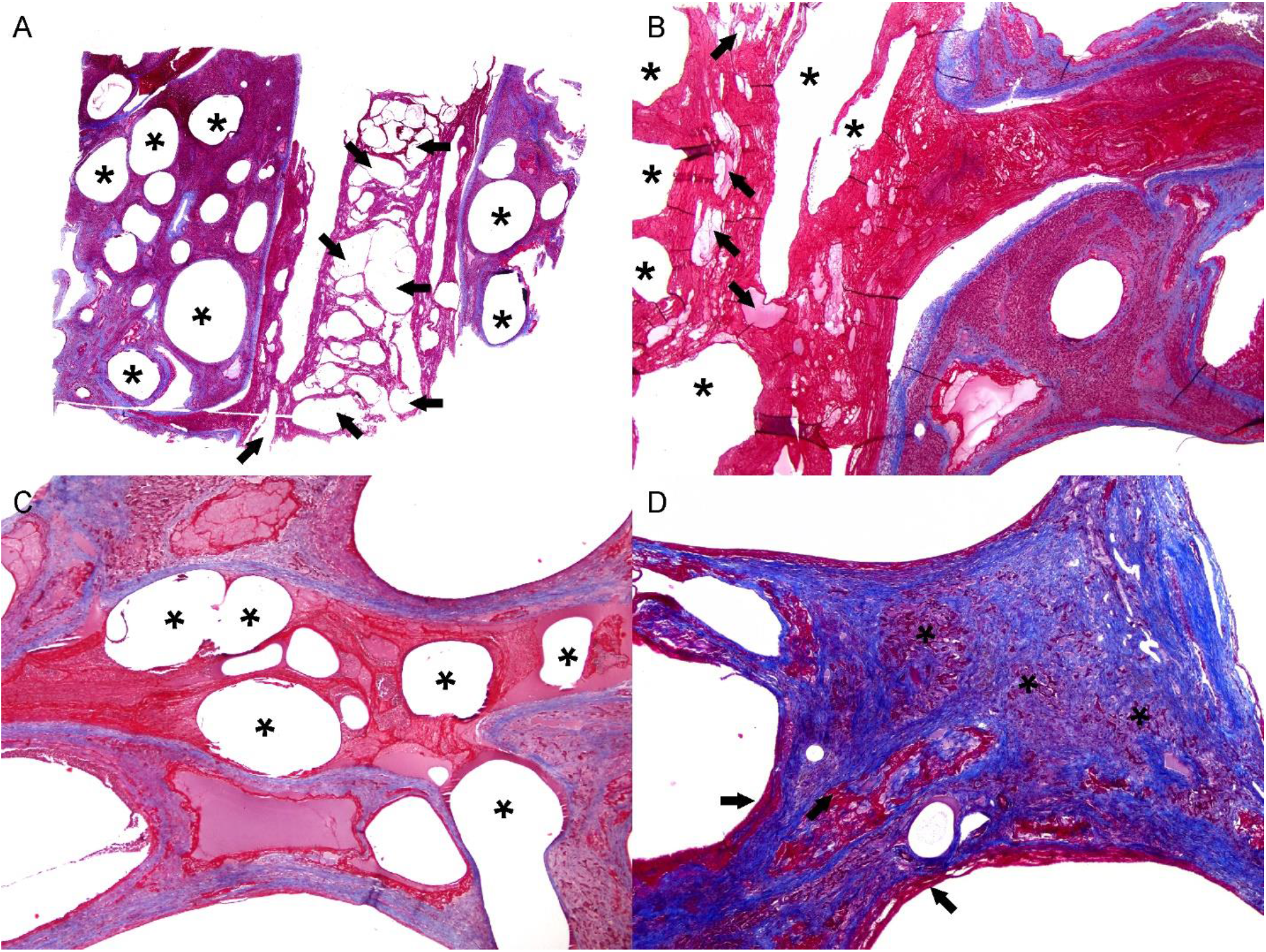
Histochemical features of cystic liver lesions in dolphins. A) Case 4. low magnification of moderate caliber hepatic vein with an obliterative thrombus. The latter contains numerous cystic spaces (arrows). Adjacent hepatic veins are cystically dilated (asterisks). B). Case 4. Higher magnification of obliterative fibrin thrombus in shown in (A). Note large organizing and branching fibrin layering and strands admixed with small pockets with serum protein (arrows) and cystic spaces (asterisks). C) Case 4. Additional poorly organized fibrin thrombi with cystic spaces (gas bubbles; asterisks) expanding serum protein, erythrocytes and fibrin. D) Case 4. Representative chronic cystic liver lesion characterized by fibrous connective tissue (blue), minimal marginal fibrin (arrows), and atrophic hepatic cords (asterisks). Trichrome stain.

**Figure 4.**
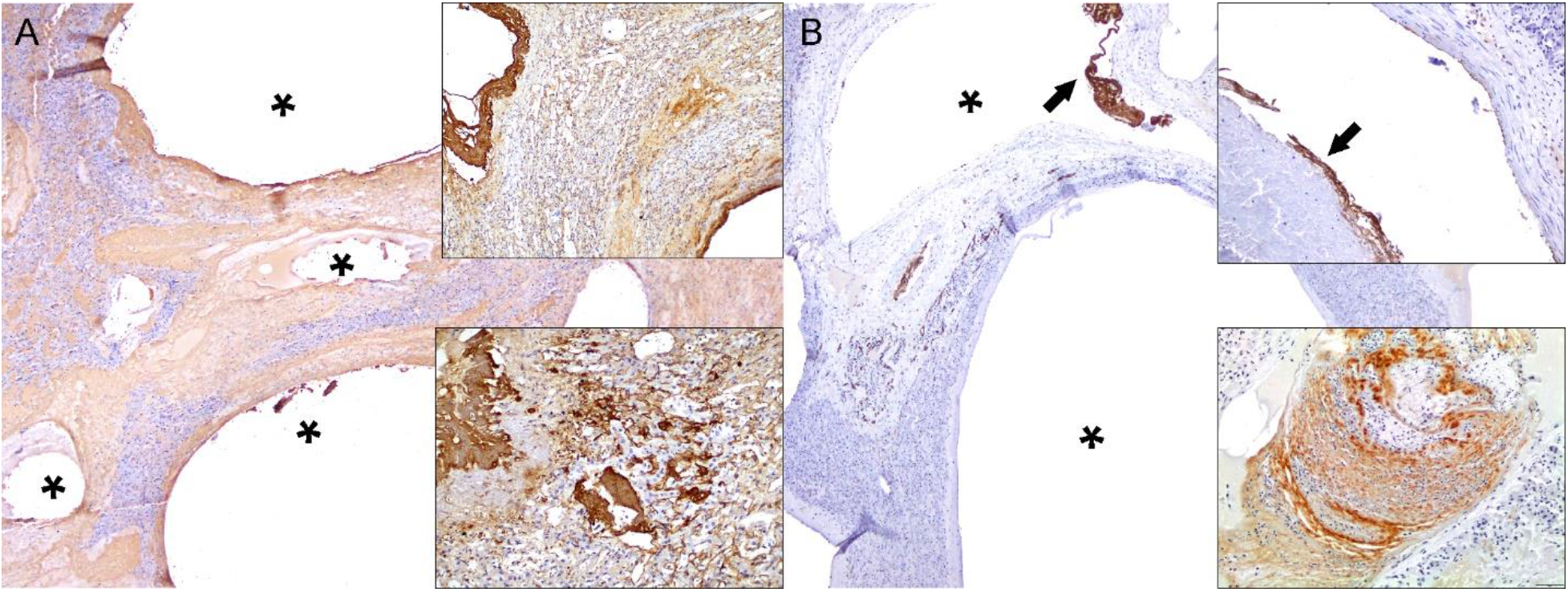
Immunohistochemical features of cystic liver lesions in dolphins. A) Case 4. Fibrinogen. Fibrinogen labeled intravascular extracellular fibrillary material surrounding intravenous cystic spaces (asterisks) as well as non-organized and organized fibrin thrombi. Right upper and lower insets: detail of intravenous and sinusoidal fibrinogen labeling. B) Case 4. Factor VIII is expressed in endothelial cells and some fibrin (arrow and right upper inset) clusters lining hepatic venous cystic dilatations (asterisks). Right lower inset: Partially organized fibrinous thrombus exhibits Factor VIII labeling.

Other relevant histopathologic findings in these animals were as follows. Case 1 had necroulcerative and suppurative gastritis with *P. gastrophilus eggs* and bacteria (pyloric chamber); chronic pancreatic ductitis/periductitis with fibrosis and intralesional trematode adults and eggs; pulmonary nematodiasis; mesangiocapillary glomerulopathy with tubular proteinosis; focal lymphocytic perivascular cuffing in cerebrum and *Nasitrema* sp. eggs associated with cranial nerves, as well as encephalic hemorrhage, edema and perivascular astrocytosis; and left pyogranulomatous orchitis with bacteria, thrombosis and fibrosis. Case 2 had necroulcerative and suppurative gastritis with perforation, fistula, and *P. gastrophilus* eggs and bacteria; peritonitis; and splenic and skeletal muscle intravascular bacteria. Case 3 had necroulcerative and suppurative gastritis with perforation and *P. gastrophilus* eggs and bacteria; severe pulmonary edema; granulomatous prostatitis; and minimal, focal lymphoplasmacytic encephalitis with lymphocytic cuffs, gliosis and satellitosis. Detailed pathologic findings in these dolphins are recorded in Supplemental table 1.

Successful IHC cross-reactivity was observed for all markers employed. Fibrinogen consistently labeled intravascular extracellular eosinophilic fibrillary material as well as non-organized and organized fibrin thrombi (Figure 3A). F-VIII was consistently expressed in endothelial cells lining hepatic arteries and veins. Rare preserved endothelial cells within early CLL were highlighted by F-VIII; some non-organized and organized thrombi expressed F-VIII (Figure 3B). The arteriolar smooth muscle cells/pericytes and vascular endothelium of the liver exhibited immunolabeling for vimentin. None of the cystic lesions had evidence of an epithelial lining based on histologic features and negativity for AE1/AE3, CK5/8 and CK8/18 cytokeratins.

### Molecular analysis

All animals tested negative for CeMV, herpesvirus and *Bartonella*.

### Gas analysis

The gas composition of both livers was very similar, and it was mainly composed of nitrogen (91.3±1.1 μmol %), with some carbon dioxide (4.7±0.4 μmol %) and oxygen (4.0±1.4 μmol %) (Table 2).

**Table 2.** Gas composition analysis results in dolphins included in this study.

| Case | Gas composition |
| --- | --- |
| 1 | Intestinal gas: 60.7% CO<sub>2</sub>; 39.3% H<sub>2</sub> |
| 2 | Not performed |
| 3 | Cystic liver lesions: 83.7 +/- 2.1 % N<sub>2</sub>; 2.7 +/- 0.5 % O<sub>2</sub>; 13.5 +/- 2.4 % CO<sub>2</sub> |
| 4 | Cystic liver lesions: 85.2 +/- 2.2 % N<sub>2</sub>; 4.8 +/- 2.3% O<sub>2</sub>; 10.1 +/-0.3 % CO<sub>2</sub> |

### Imaging and liver cast analysis

CT-scan and liver vasculature corrosion cast model analyses enabled visualization and discernment of the vastly rich vasculature of the liver in two control striped dolphins, one each (Supplementary Figures 1 and 2). CLL were consistently observed in the upper right and lower quadrant of the right liver lobe (4/4) while only one dolphin had lesions on the left liver lobe. The main vessels affected were (in decreasing order), the large right hepatic veins and sinuses, and right medium and smaller caliber hepatic veins. Furthermore, there was a clear dorso-caudal to cranio-ventral progression of CLL.

### Bacteriologic analyses

Bacterial isolates are recorded in Supplemental table 3. Case 1 (brain, spleen, liver, kidney, skeletal muscle, lung) yielded no bacterial growth. Case 2 yielded *Enterococcus faecalis* pure culture from spleen, liver, kidney, adrenal gland, prescapular and mesenteric lymph nodes. Cases 3 yielded *Enterococcus thailandicus* and *E. faecium* from brain; *Stenotrophomonas maltophilia* and *S. nitritireducens* from liver; *E. thailandicus* from prescapular, pulmonary and mesenteric lymph nodes; and *E. thailandicus* and *E. hirae* from oral cavity, esophagus, and heart.

## Discussion

Based on these results, we infer that CLL in striped dolphins result from the combination of a pre-existing or concomitant hepatic vascular disorder (HVD) superimposed and/or may be exacerbated by *ex novo*, *in situ* formed or circulating gas (from decompression phenomena) embolic bubbles resulting in hepatic cysts mainly containing nitrogen^8^. These stranded dolphins were either not associated temporally nor spatially with any primary anthropic acoustic activities. We propose this condition to be named preliminarily “cetacean polycystic liver pathology” (CPLP). The latter term is proposed until *“in vivo”* diagnosis and clinical prognosis allow for a better characterization of the condition, at which point cetacean polycystic disease could be more appropriate.

The gross, histologic, histochemical and immunohistochemical findings of CLL in this study ratified previous gross and histologic observations^1,2,10^ and allowed for identification of the progressive or chronic-active nature of the disease process. The histologic features of CLL were similar in all cases and suggested an ordered progression that included formation of gas-filled cystic spaces within the matrix of thrombosed hepatic veins into their intraparenchymal branches and hepatic parenchyma. Specifically, “early” gas-filled cystic cavities arose within thrombosed and obliterated segments of large, medium and small caliber intraparenchymal hepatic veins; smaller cystic spaces forming peliotic foci within lobules affecting a variable extension of sinusoids were also common. In these early stages, cystic spaces had abundant serum protein, fibrin with varying levels of organization, erythrocytes, and rare leukocytes. The endothelial lining in these early lesions was occasionally identifiable and immunolabeled with FVIII. As these lesions progressed, cystic spaces had wider non-stained (gas-filled) areas, thinner peripheral rims of fibrin, and progressive deposition of fibrous connective tissue, as highlighted by trichrome and picrosirius red stains. Early and advanced lesions coexisted and were occasionally associated with fatal large infarcts; these were particularly prominent in case 4. The more advanced lesions lacked endothelial cells. Fibrinolysis likely accounted for lack of fibrin in the most chronic lesions. Fibrous deposits included original perivascular intermediate filaments plus subsequent collagen deposition. F-VIII and fibrinogen IHC revealed endothelial damage followed by remodeling of vessel walls and platelet and fibrin distribution through early and advanced lesions. Cytokeratin immunomarkers discarded a biliary origin for CLL, therefore, these lesions are clearly different from developmental and acquired biliary cystic diseases as reported in humans and other animal species.

Noteworthy, none of these dolphins had evidence of systemic acute gas embolism, and no cystic lesions were observed in the spleen, kidney or any other tissue examined. Furthermore, based on parallel diagnostic imaging and liver vasculature corrosion model of control liver of striped dolphins, it was evident that CLL consistently involved the upper right and lower quadrant of the right liver lobe, corresponding to the main hepatic veins in these regions. Only one dolphin had lesions on the left liver lobe coinciding with the most severe presentation recorded in this set of animals. Furthermore, there was a clear dorso-caudal to cranio-ventral progression of CLL, which is in agreement with progressive reduction of caliber of hepatic veins.

In humans, HVD may be classified according to the type of blood vessels involved, namely portal vein, hepatic artery, sinusoids, and hepatic veins^25^. The obliterative thromboembolic lesions observed in these dolphins were primarily centered on large hepatic veins including the typical sinuses of cetaceans, as well as medium and small caliber hepatic veins, recapitulating the anatomic distribution and histopathologic features of Budd–Chiari syndrome (BCS)^25^. BCS denotes hepatic venous outflow obstruction, regardless the level or the mechanism of obstruction. Furthermore, BCS is classified as ‘primary’ when it is related mainly to venous disease, such as thrombosis or phlebitis, and ‘secondary’ when it is caused by venous compression or invasion by a lesion originating outside the veins, such as an abscess or a tumor.^25^ In these striped dolphins, it is reasonable to note that a secondary phenomenon, namely, severe trematodal cholangiohepatitis resulted in localized vascular involvement and eventual phlebothrombosis. The latter might have served as matrix for entrapment of circulating gas bubbles *in situ*, or for *de novo* gas bubble formation. Portal veins were rarely affected; hepatic arteries were largely unaffected. Comparatively, human BCS cases lack the herein proposed critical secondary etiopathogenic factor, namely intravascular gas bubbles. A putrefactive phenomenon, portal embolism from intestinal gas or septic enteric gas-forming process^1,2^ were reasonably excluded on the basis of the gas profile in cases 3 and 4 as well as lack of grossly evident mesenteric gas. Notably, no pyogenic or neutrophilic inflammation (other than the argued predisposing cause) was observed associated with CLL.

The exact etiopathogenesis(es) of CLL in cetaceans warrants further investigation. In humans, BCS has been ascribed to a plethora of causes which may be further subdivided in a) hypercoagulable states, b) stasis or mass lesions, c) vascular injury, d) surgical manipulation, and e) uncertain mechanisms^25^. The potential for overlapping features in many of these conditions may be an indication of the intricacy yet interconnection and common involvement of different vascular segments in extensive hepatic lesions. Once organizing fibrin thrombi/thromboemboli within hepatic veins are stablished, previously circulating and entrapped gas bubbles and/or *de novo*, *in situ* gas bubbles may increase in number, resulting in increased resistance to blood flow, congestion, secondarily peliotic (phlebectatic, telangiectatic) lesions as well as reciprocal venous thrombosis. In the end, a vicious pathologic cycle may ensue leading to severe, coalescing, and extensive CLL as it has been observed in these striped dolphin livers and other species^2^.

Furthermore, our results indicate CLL occurred in stranded cetaceans with poor or emaciated body condition and multiorgan lesions. It is possible that as CLL progresses it may cause congestive hepatopathy, portal hypertension, multisystemic hypertension and impairment of general host homeostasis, resulting in a progressive debilitating multiorgan disease which could potentially favor and/or aggravate concomitant disease processes. Our findings lend support to this interpretation. The current data also suggest that individual factors, such as species and individual health status, may play a role as predisposing factors in CLL. Consistent concomitant pathologies in these dolphins were moderate to severe gastric trematodiasis by *P. gastrophilus* and pancreatic trematodiasis by *Brachycladiidae*. In case 2, gastric and pancreatic trematodiasis related to severe hepatic abscess, peritoneal effusion and peritonitis, and likely resulted in septic peritonitis and bacteremia. Cases 1 and 3 had evidence of minimal inflammatory foci in the encephalon; the exact etiology remains uncertain, however, nasitremiasis could have played a role in case 1. All these cases were PCR-negative for CeMV, HV and *Brucella*, known causes of nervous and multiorgan inflammatory disease. Most other lesions observed in these animals were the result of chronic endoparasitism (e.g., pulmonary nematodiasis) or concomitant bacterial infections (e.g., orchitis and peritesticular abscess). Bacterial analyses results indicated *Enterococcus faecalis* played a role in peritonitis and septicemia in case 2. The significance of various bacteria identified in case 3 is uncertain.

Interestingly, none of the 100+ beaked whales fully examined by our team to date have had any evidence of CLL. Also noteworthy, none of the examined beaked whales that stranded in association temporally and spatially with sonar exposure had CLL; instead, those animals presented acute systemic gas embolism findings consistent with DCS^26,5,9^. Furthermore, our hypothesis of focal/multifocal vasculopathy and thrombosis followed by intravascular gas-bubble entrapment and expansion resulting in localized necrosis, could be a suitable fit to explain development of dysbaric osteoneocrosis, yet another highly controversial pathologic finding in sperm whales allegedly associated with DCS^27^.

In conclusion, our results ratified the occurrence of *in vivo* decompression phenomena and strongly suggest that CLL are the result of the combination of pre-existing or concomitant hepatic vascular disorder superimposed and exacerbated by gas bubbles (gas embolism). CPLP clearly differs from acute systemic gas embolism reported in stranded beaked whales linked to MFAS. Specifically, in these dolphins, CPLP was the result of hepatic vascular disorder and phlebothrombosis plus entrapment of decompressing formed intravascular circulating gas bubbles. CPLP invariably involved dolphins with a poor body condition and concomitant chronic multiorgan disease processes, including severe hepatobiliary trematodiasis. CPLP should be interpreted as a chronic, progressive hepatic vascular disease process with potential to result in significant low individual morbidity and mortality also in striped dolphins.

## Methods

### Animals investigated

The Canarian Marine Mammal Stranding Network has performed over 1,200 necropsies in cetaceans stranded, including 172 striped dolphins (*Stenella coeruleoalba*). Out of these, 4 (2%) striped dolphins have shown CLL similar or identical to those previously reported in Risso’s dolphins, common dolphins, and harbor porpoises^1,2^. This study focused on those four striped dolphins. Biologic and stranding epidemiology data for each individual are recorded in Table 1. Age category was based on total body length and gonadal development, including fetus/neonate/calf, juvenile/subadult, and adult^11,12,13^. The nutritional status (syn. body condition) was subjectively classified into good, moderate, poor, and emaciated according to anatomic parameters such as the osseous prominence of the spinous and transverse vertebral processes and ribs, the mass of the epaxial musculature, and the amount of fat deposits, taking into account the species and the age of the animal^11,12^. Carcasses were classified as very fresh, fresh, moderate autolysis, advanced autolysis or very advanced autolysis^11,12,13,14^.

Required permission for the management of stranded cetaceans was issued by the environmental department of the Canary Islands’ Government and the Spanish Ministry of Environment. No experiments were performed on live animals.

### Pathologic examinations

Necropsies followed standardized protocols^11,12,13,14^. Representative samples of skin, *longissimus dorsi* and *rectus abdominis* muscles, peritoneum, diaphragm, central nervous system, eye, pterygoid sac, tympanoperiotic complexes, tongue, oral mucosa, pharyngeal and laryngeal tonsils, esophagus, stomach, small and large intestine, liver, pancreas, trachea, lung, heart, aorta, kidney, ureter, urinary bladder, urethra, lymph nodes, spleen, testicle, penis, prepuce, ovary, uterus, vagina and vulva, were collected and fixed in 10% neutral buffered formalin. All these tissues were processed routinely, embedded in paraffin-wax and 5 μm-thick sections were stained with hematoxylin and eosin (H&E) for microscopic analysis. Special histochemical techniques (4-10 μm-thick sections) to better characterize microscopic features of CLL included periodic acid-Schiff, Masson’s trichrome, Picrosirius red, Perl’s and Warthin-Starry.

### Immunohistochemical analysis

We employed a set of primary antibodies for immunohistochemical (IHC) analysis to better characterize CLL. Primary antibodies utilized were factor (F)-VIII (syn. Von Willebrand factor), for endothelial cells and platelets; fibrinogen, as a proxy of fibrinogen/fibrin (thrombosis); pancytokeratin AE1/AE3 and cytokeratins 5, 8 and 18 for bile duct epithelium; and vimentin as general marker of intermediate filament for mesenchymal cells. Published methodologies for IHC analyses were followed^15,16^. Detailed information on IHC methodology is recorded in Supplemental table 2. Negative controls consisted of serial tissue sections in which primary antibodies were substituted by non-immune homologous serum. Positive controls included internal and external striped dolphin liver with its normal constituents.

### Polymerase chain reaction analyses

At necropsy, selected samples were collected for virologic analyses and stored frozen at −80°C until processing for molecular virology testing. Approximately 0.5 gr of fresh-frozen tissue sample from each animal was mechanically macerated in lysis buffer and subsequently centrifuged. DNA/RNA extraction was carried out from each 300 μL macerated sample by pressure filtration method, using a QuickGene R Mini 80 nucleic acid isolation instrument, using the DNA Tissue Kit S (QuickGene, Kurabo, Japan) according to the manufacturer’s instructions with modifications: RNA carrier (Applied Biosystems, Thermo Fisher Scientific Waltham, Massachusetts, USA) was added during the lysis step^17^. Herpesvirus DNA was detected by conventional nested PCR using degenerate primers designed to amplify a region of the DNA polymerase gene^18^. Molecular detection of cetacean morbillivirus (CeMV) was performed by one or more of four different PCR methods: one-step RT-PCR of a 426-bp conserved region of the phosphoprotein (P) gene^19^, one-step real-time RT-PCR that detect the most common CeMV strains known to circulate in the Atlantic Ocean targeting the P gene^20^, and RT-PCR using nested primers targeting the P gene^21^, and one-step real-time RT-PCR to detect sequences in a conserved region (192 bp) of the fusion protein (F) gene^17^. Molecular detection of *Bartonella* spp. was additionally carried out by means of a real-time PCR assay using SYBR® Green, as previously described^22^.

### Gas sampling and analysis

Gas was sampled from CLL of cases 3 and 4 and intestine of case 1. To prevent atmospheric air contamination, the livers were submerged in distilled water and gas was sampled using an aspirometer (ES1076326) following Bernaldo de Quirós *et al*. (2011)^8^. Gas samples were collected and stored in 5-mL additive-free glass vacutainers® (BD Vacutainer® Z. ref: 367624) and analyzed by gas chromatography within two weeks of sample collection. Gas samples were extracted from the vacutainers and injected into the gas chromatograph (Varian 450-GC) using glass block-pressure syringes (Supelco A-2 series). Gas samples were run for 15 min at 45**°**C and with a fixed pressure of 13.1 psi through a Varian 450-GC column and a thermal-conductivity detector as well as a flame-ionization detector using Helium as the carrier gas^8^. The results were expressed as the average ± the standard deviation in μmol %.

### Computed tomography

A normal liver of an adult striped dolphin (for comparative purposes) was scanned with a 16-slice helical CT scanner (Toshiba Astelion, Toshiba Medical System, Madrid, Spain) under the following settings: 16 detector arrays; type of acquisition: helical; thickness: 1 mm; image reconstruction interval: 0,8 mm; pitch: 1; tube rotation time: 0,75; mA 250; Kv: 120; FOV: 512X512; Matrix dimensions: 1204 x 845; WW 500/WL 505. Previously, the liver was dissected; the caudal vena cava and the portal veins were isolated and cannulated separately. These vessels were flushed with tap water and then a radiocontrast agent (Iomeron 300©) mixed with saline solution was injected in both veins. The resulting DICOM images were analyzed; a soft tissue window setting (WW 500/WL 50) was applied making a rendering of the internal volume and thus achieving a 3D model of both venous systems, portal and hepatic.

### Liver cast analysis

For detailed assessment of the hepatic vasculature, we performed a corrosion cast on a control striped dolphin (comparative purposes) following published methodologies^23,24^. At necropsy, the caudal vena cava, portal vein, hepatic artery and choledochal duct were carefully dissected, isolated and flushed with tap water. Then, the liver was carefully dissected and detached from the supporting vessels while preserving the capsule and its ligaments (it was extracted with part of the diaphragm and the lesser omentum). Once removed, the liver was carefully washed. *Ex situ*, the common bile duct, hepatic artery, portal vein, and caudal vena cava were catheterized at the dorsal edge of the liver. The left branch of the caudal vena cava was clamped. Following, blue, white, red and yellow epoxy (Biodur©) was injected into the venous and arterial vessels and the choledochal duct, respectively. The liver was stored at room temperature for 24h and then at 4°C for 24h. After that, a corrosion process was applied over the submerged liver with a 10% sodium hydroxide solution (NaOH) for two weeks. The soft tissue debris were removed manually.

### Bacteriologic analyses

Fresh tissue samples (skin, muscle, lung, prescapular, pulmonary, mediastinal and mesenteric lymph nodes, liver, intestine, kidney, spleen, brain) collected routinely during necropsy, were frozen (−80°C) and selectively submitted for bacteriologic analysis. These included routine culture and surface plating on routine media, *e.g*., Columbia blood agar and preliminary identification of isolates via API® system (API® 20E, API® Rapid 20E, API® Staph, API® 20 Strep, API® Coryne, API® 20A). PCR targeting the 16S rRNA gene coupled with pulsed-field gel electrophoresis were performed on selected isolates.

## Acknowledgements

This study was supported by the National Project PGC2018-101226-B-I00 and the Canary Islands Government, which has funded and provided support to the stranding network. The authors would like to thank to Xavier Lekuve at Biscay Bay Environmental Biospecimen Bank (BBEBB) Estación Marina Plentzia (PiE-UPV/EHU) and Leire Ruiz (AMBAR Elkartea), Secac, Marisa Tejedor and Canarias Conservacion for their collaboration.

## Author contributions

A.F. contributed to the conception or design of the work and interpretation of data. AF, JDD, YBQ, MA, EV, MR, FC, BM, AEM, PH, MA, AIV, CSS, MJC, OQ, AC, GDG have provided necropsy work and laboratory results, discussion, drafting and review the work at different stages of this 20 years’ work. PDJ provided the original work previously published in Nature and to whom this article is dedicated.

## Data availability

The authors declare that the data supporting the findings of this study are available within the paper and its supplementary information files.

---

**Supplementary Figure 1.**
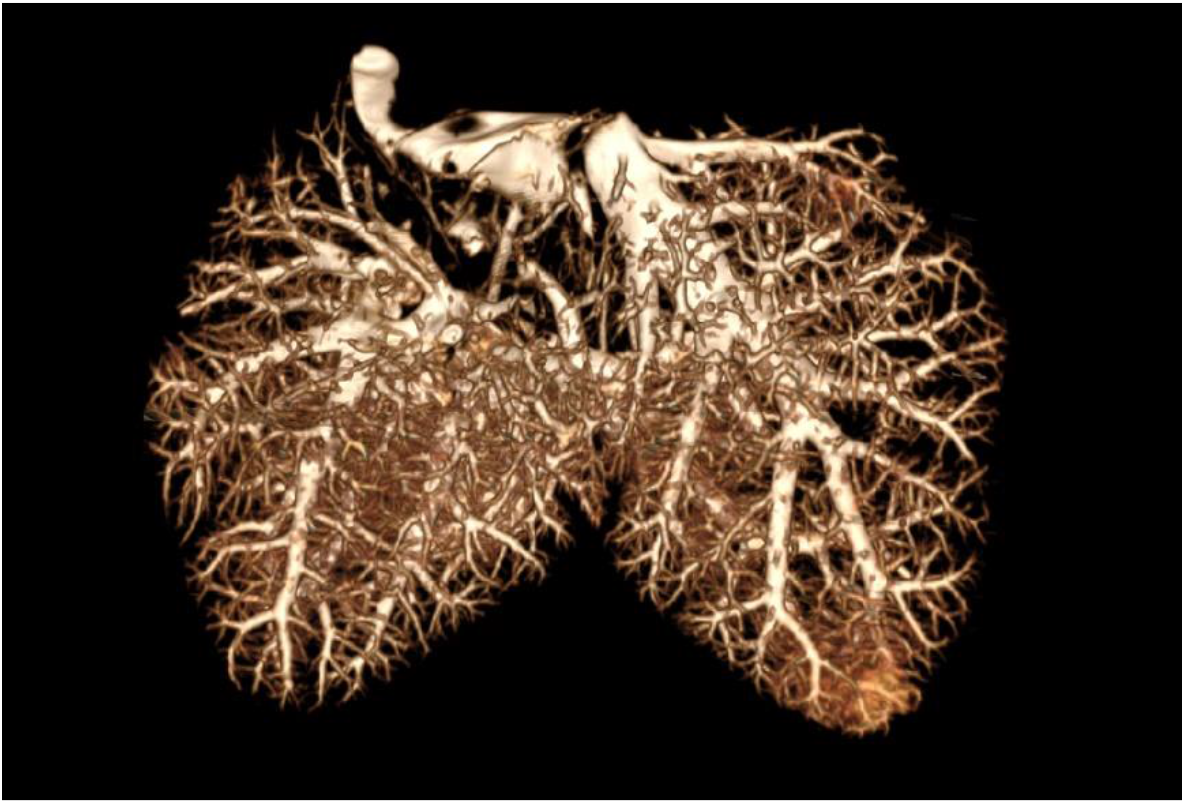
CT-scan rendering of liver vasculature in control striped dolphin.

**Supplementary Figure 2.**
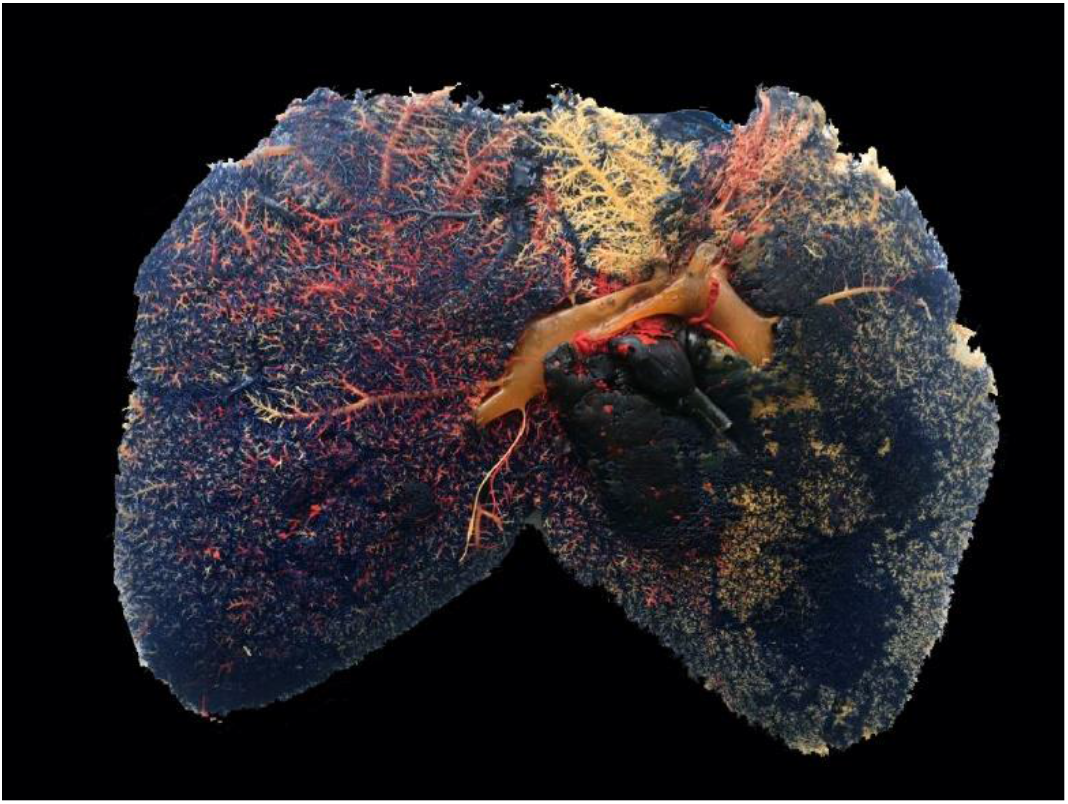
Liver vasculature corrosion cast model in control striped dolphin.

**Supplementary table 1.** Detailed gross and histopathologic findings in the animals included in this study.

| Case | Gross findings | Histologic findings |
| --- | --- | --- |
| 1 | **Skin:** Numerous tooth rake marks throughout the bodily surface; Stellate scars (*Isistius* sp. bite wounds) on the chest; Granulomatous panniculitis with yellow to dark brown foci; Bilateral thoracolumbar edema; Severe subcutaneous infection by *P. delphini* merocercoids in anogenital region and peduncle.<br>**Left testicle:** Focal 31 x 23 cm, peritesticular/ epididymal abscess spanning from the testicle into the cranial abdomen displacing the small intestine and compressing the colon. Upon section, 2.5 L of purulent exudate oozed from the abscess; Diffuse testicular atrophy.<br>**Oral cavity:** Multiple teeth missing.<br>**Keratinized stomach:** Severe anisakiasis; Small number of squid beaks and otoliths.<br>**Glandular and pyloric stomachs:** Ulcerative gastritis.<br>**Liver (non-cavitary lesions):** Severe infection by Brachycladiidae trematodes with bile ducts and pyogranulomas. Diffuse hepatic congestion.<br>**Peritoneal cavity:** Severe infection by *M. grimaldi* merocercoids.<br>**Lung:** Diffuse pulmonary edema and congestion; Bilateral intraparenchymal granulomas and mild intrabronchial nematodes.<br>**Kidney:** Bilateral diffuse congestion<br>**Ureter:** Mild bilateral ureterectasis (hydroureter).<br>**Pulmonary and prescapular lymph nodes:** lymphadenomegaly with intralesional nematodes. | **Skin:** Granulomatous dermatitis and panniculitis with phagocytized and extracellular yellow globules.<br>**Skeletal muscle:** Acute myofiber degeneration and necrosis with phagocytosis and occasional regeneration; Myocyte atrophy.<br>**Gastric chambers:** Lymphoplasmacytic and eosinophilic gastritis with *P. gastrophilus* eggs, superficial bacteria and fibrosis (glandular chamber); Necroulcerative and suppurative gastritis with *P. gastrophilus* eggs and bacteria (pyloric chamber).<br>**Intestine:** Lymphoplasmacytic and eosinophilic enteritis.<br>**Pancreas:** Chronic ductitis/periductitis with fibrosis and intralesional trematode adults and eggs.<br>**Liver (non-cavitary lesions):** Lymphoplasmacytic and neutrophilic cholangiohepatitis with necrosis, bile duct hyperplasia, fibrosis, hemorrhage and hemosiderosis (intravascular and intralesional bacteria); Severe, multifocal, suppurative cholangiohepatitis with biliary and hepatic necrosis, trematode adults and eggs, and bacteria.<br>**Lung:** Lymphoplasmacytic bronchiointerstitial pneumonia with pyogranulomas, nematodes, fibrosis, angiomatosis, hemosiderosis; Atelectasis and emphysema; Bronchial/-olar mineralization.<br>**Heart:** Multifocal, acute cardiomyocyte degeneration; Multifocal arterial tunica media hypertrophy/hyperplasia.<br>**Kidney:** Mesangiocapillary glomerulopathy with tubular proteinosis and tubuloe epithelial pigment; Tubular mineralization.<br>**Urinary bladder:** Lymphoplasmacytic and histiocytic serositis with hemorrhage, hemosiderosis and focal *M. grimaldi* merocercoid.<br>**Thyroid:** Perithyroid hemorrhage and thrombosis; Interstitial fibrosis.<br>**Adrenal gland:** Lymphoplasmacytic adrenalitis and fibrosis.<br>**Cerebrum, Cerebellum:** Hemorrhage, edema and perivascular astroglyosis; Minimal, focal lymphocytic perivascular cuffing; Choroid plexus hyalinization; *Nasitrema* sp. eggs associated with cranial nerves.<br>**Pulmonary, Prescapular, lymph node:** Sinus histiocytosis and neutrophilia with intravascular and sinus bacteria, sinus yellow globules, and gas/fat sinus dilatations; Lymphoid depletion.<br>**Spleen:** Lymphoid depletion and sinus histiocytosis and hemosiderosis; Extramedullary hematopoiesis.<br>**Right testicle:** Lymphoplasmacytic interstitial orchitis and epididymal hemorrhage.<br>**Left testicle:** Pyogranulomatous orchitis with bacteria, thrombosis and fibrosis. |
| 2 | **Skin:** Numerous tooth rakes throughout the bodily surface; Irregular 20-cm-long irregular healed wound on ventral abdomen and stellate pigmented wounds; Mild subcutaneous infection by *P. delphini* merocercoids in anogenital region and peduncle.<br>**Cephalic skeleton:** Rostral fractures with soft tissue loss.<br>**Peritoneal cavity:** Moderate amount of red exudate within the peritoneal cavity associated with serosal villous projections (chronic serositis); Moderate infection by *M. grimaldi* merocercoids.<br>**Liver:** Focally extensive 18 cm-diameter abscess at distal aspect and parietal surface of liver, firmly adhered to the adjacent pyloric compartment. On cut surface, there is necrosis and fibrinopurulent exudate, fibrosis, adult flukes and larvae | **Skin:** Lymphohistiocytic dermatitis and panniculitis.<br>**Glandular stomach:** Chronic lymphoplasmacytic gastritis with lymphonodular hyperplasia<br>**Pyloric stomach:** Necroulcerative and suppurative gastritis with perforation, fistula and *P. gastrophilus* eggs and bacteria.<br>**Liver (non-cavitary lesions):** Pyogranulomatous cholangiohepatitis with trematode adults and eggs, bacteria, necrosis, fibrosis and neovascularization.<br>**Lung:** Intravascular and intrabronchiolar cystic dilatations; Mixed intrabronchiolar exudate; Bronchial/-olar mineralization.<br>**Mediastinal, mesenteric lymph nodes:** Intrasinus and intravascular cystic spaces.<br>**Heart:** Leukocytosis; Acute cardiomyocyte degeneration; Rare perivascular and interstitial neutrophils; Interstitial edema.<br>**Spleen:** Intravascular bacteria<br>**Skeletal muscle:** Intravascular/interstitial bacteria. Rare acute myocyte degeneration; Rare myocyte regeneration. |
|  | (*Campula* sp. and *Pholleteer gastrophilus*, presumably), and otoliths.<br>**Pyloric chamber:** Firmly adhered to the hepatic abscess. Upon dissection, a 3 cm-diameter fistulous track communicated the pyloric compartment and liver parenchyma (focal transmural pyogranulomatous gastritis with fistulae, necrosis, intralesional adult flukes (*P. gastrophilus*) and adjacent liver fistula and adhesion; Mild anisakiasis.<br>**Pancreas:** Numerous trematode adults and larvae (*Campula* sp.) in hepatopancreatic ducts and hepatic hilum.<br>**Cephalic skeleton:** Intermandibular symphyseal fracture with tissue loss.<br>**Larynx:** Mild infection by 1.5-2.5 x 0.05 cm nematodes.<br>**Lung:** Diffuse atelectasis with 5-17 x 2-6 cm subpleural bullae on left lung. Focal 5.5 cm subpleural hemorrhage; Marginal atelectasis and multifocal subpleural emphysema; Mild intrabronchial nematodal infection identical to those in larynx.<br>**Peritoneal cavity:** Vascular dilatation in mesentery, gastrosplenic and perirenal regions; mild in coronary. |  |
| 3 | **Skin:** Multifocal pigmented circular cutaneous lesions; Moderate, multifocal intra-specific interaction marks (tooth rakes) in peduncle; Partial amputation of dorsal fin with complete scarring; Live-stranding related linear erosions, abrasions and ulcerations throughout the ventrolateral bodily surface; Erosions and ulcers along the cranial edge of pectoral fins, eye and beak; Focal 1.5 cm in diameter, circular, depressed focus with red margins dorsally to right flipper; Moderate infection by *P. delphini* in ventrocaudal region; Serous atrophy of fat (cervical region).<br>**Oral cavity:** Marked dental wear and multiple teeth absence; Chronic gingivostomatitis.<br>**Tongue:** Chronic glossitis with hyperplastic plaque-like foci, erosions and ulcers.<br>**Glandular and pyloric stomachs:** Proliferative and granulomatous gastritis with *P. gastrophilus*; Three squid beaks.<br>**Trachea, bronchi, lungs:** Marked pulmonary edema.<br>**Right lung:** Atelectasis of left lung; Emphysema of right lung; Bilateral pulmonary nematodiasis.<br>**Heart:** Hemorrhage in aortic valves.<br>**Peritoneal cavity:** Moderate infection by *M. grimaldii* in mesentery and peritoneum.<br>**Kidney:** Bilateral nephrolithiasis.<br>**Prostate:** Granulomatous prostatitis. | **Skin:** Proliferative dermatitis with intracytoplasmic inclusions (compatible with *Poxvirus*); Neutrophilic and ulcerative dermatitis and panniculitis with bacteria.<br>**Cutaneous trunci muscle:** Suppurative myositis.<br>**Skeletal muscle:** Myocyte atrophy and regeneration.<br>**Diaphragm:** Interstitial edema.<br>**Lung:** Severe alveolar edema, congestion and hemorrhage; Atelectasis and emphysema; Bronchial/olar mineralization; Lymphoplasmacytic and neutrophilic interstitial pneumonia.<br>**Oral cavity:** Marked dental wear; Chronic gingivostomatitis.<br>**Tongue:** Multifocal proliferative and ulcerative glossitis.<br>**Esophagus:** Lymphoplasmacytic esophagitis.<br>**Glandular and Pyloric stomachs:** Necroulcerative and suppurative gastritis with perforation and *P. gastrophilus* eggs and bacteria.<br>**Intestine:** Eosinophilic enteritis.<br>**Liver (non-cavitary lesions):** Chronic cholangiohepatitis with fibrosis; Marked cholestasis.<br>**Kidney:** Lymphoplasmacytic interstitial nephritis with fibrosis; Tubular mineralization and nephrolithiasis.<br>**Prostate:** Granulomatous prostatitis.<br>**Prescapular, Mesenteric lymph node:** Sinus histiocytosis and hemosiderosis; Lymphoid depletion; Eosinophilic lymphadenitis.<br>**Spinal cord, Cranial nerves:** Congestion and leukocytosis.<br>**Cerebrum:** Minimal, focal lymphoplasmacytic encephalitis with lymphocytic cuffs, gliosis and satellitosis. |
| 4 | Not recorded | Not recorded |

**Supplementary table 2.** Methodology for immunohistochemical analyses employed.

| Marker | Clonality | Host | Source | Antigen retrieval | Dilution |
| --- | --- | --- | --- | --- | --- |
| **AE1/AE3** | Monoclonal | Mouse | Biocare Medical | 10% pronase | 1:2,000 |
| **CK5/8** | Monoclonal | Mouse | Euro-Diagnostica | 10% pronase | 1:20 |
| **CK8/18** | Monoclonal | Mouse | Euro-Diagnostica | Citrate buffer | 1:20 |
| **Vimentin** | Monoclonal | Mouse | Dako | Citrate buffer | 1:100 |
| **Fibrinogen** | Polyclonal | Rabbit | Abcam | Citrate buffer | 1:50 |
| **Factor VIII** | Polyclonal | Rabbit | Thermo Fisher Scientific | 0.1% trypsin | 1:100 |

**Supplementary table 3.** Microbiologic results in the animals included in this study.

| Case | Tissues | Bacteriology |
| --- | --- | --- |
| 1 | Brain, spleen, liver, kidney, skeletal muscle, lung | No growth |
| 2 | Spleen, liver, kidney, adrenal gland, prescapular and mesenteric lymph nodes | *Enterococcus faecalis* |
| 3 | Brain | *Enterococcus thailandicus*, *E. faecium* |
|  | Liver | *Stenotrophomonas maltophilia*, *S. nitritireducens* |
|  | Prescapular, pulmonary and mesenteric lymph nodes | *Enterococcus thailandicus* |
|  | Oral cavity, esophagus, heart | *Enterococcus thailandicus*, *E. hirae* |

